# A graph-based algorithm for RNA-seq data normalization

**DOI:** 10.1101/321471

**Authors:** Diem-Trang T. Tran, Aditya Bhaskara, Matthew Might, Balagurunathan Kuberan

## Abstract

The use of RNA-sequencing has garnered much attention in the recent years for characterizing and understanding various biological systems. However, it remains a major challenge to gain insights from a large number of RNA-seq experiments collectively, due to the normalization problem. Current normalization methods are based on assumptions that fail to hold when RNA-seq profiles become more abundant and heterogeneous. We present a normalization procedure that does not rely on these assumptions, or on prior knowledge about the reference transcripts in those conditions. This algorithm is based on a graph constructed from intrinsic correlations among RNA-seq transcripts and seeks to identify a set of densely connected vertices as references. Application of this algorithm on our benchmark data showed that it can recover the reference transcripts with high precision, thus resulting in high-quality normalization. As demonstrated on a real data set, this algorithm gives good results and is efficient enough to be applicable to real-life data.

**2012 ACM Subject Classification:** Applied computing → Computational transcriptomics, Applied computing → Bioinformatics

**Digital Object Identifier:** 10.4230/LIPIcs.WABI.2018.xxx

**Funding:** This material was based on research supported by the National Heart, Lung, and Blood Institute (NHLBI)–NIH sponsored Programs of Excellence in Glycosciences [grant number HL107152 to B.K.], and partially by NSF [CAREER grant 1350344 to M.M.]. The U.S. Government is authorized to reproduce and distribute reprints for Governmental purposes notwithstanding any copyright notation thereon.

## 1 Introduction

RNA-sequencing (RNA-seq) has become a critical tool to study biological systems [15]. The technique starts with extracting the RNA fraction of interest and preparing them for high-throughput sequencing. Sequencers typically output short reads that are then assembled or aligned to a pre-assembled genome or transcriptome, resulting in a quantity called *read count* for each transcript. Due to variations in sequencing depth (i.e. library size, the total number of read count per sample) and in relative contribution of each transcript under different conditions, these read counts need to be normalized such that the changes in their measurements, usually indicated by fold-change, accurately reflect the differences between conditions. This is often named the between-sample normalization problem, which has attracted much efforts in solving it. These solutions vary in their approaches and more importantly, their assumptions [5]. The proposed solutions so far can be grouped into two major classes, following the classification by Evans et al. [5]: normalization by distribution/testing and normalization by references^1^. Benchmark studies have come to support several methods in the first group such as TMM, DESeq due to their good performance in the differential expression (DE) analyses [3, 10]. A major caveat with these methods is the core assumption that most genes are not differentially expressed across conditions of interest. This assumption fails to hold when one needs to analyze multitude and variable conditions such as tissue types. In the later group of methods, reference-based normalization, a subset of transcripts that are stably expressed across conditions will be used to calculate the normalizing factors. The identification of these transcripts initially depended on their functional annotations and a seemingly valid rationale that housekeeping genes are universally required for critical living functions, thus should be expressed at similar levels across all different conditions. Many housekeeping genes commonly used as references turned out to vary significantly across conditions (see Huggett et al. [8] for an extensive list of such examples), leading to the favor of external references, i.e spike-in RNAs [9, 2] or automatic methods to determine internal references [1, 16]. The addition of external spike RNAs significantly increases the cost, complicates experimental processes, and is inapplicable for integrating the large number of data from different experiments and laboratories which used different spikes, or most of the times, no spike at all. Automatic identification of stably expressed genes, on the other hand, is usually based on some coefficient of variation which requires a proper normalization, therefore unavoidably suffers from a circular dependence [16].

We propose a new method of normalization that can break this circularity, without relying on any assumption about the biological conditions at hands. We show that there exist intrinsic correlations among reference transcripts that could be exploited to distinguish them from differential ones, and introduce an algorithm to discover these references. This algorithm works by modeling each transcript as a vertex in a graph, and correlation between them as edges. In this model, a set of references manifest themselves as a complete subgraph and therefore can be identified by solving a clique problem. We also show that this algorithm, with a few practical adjustments, can be finished in reasonable time and give good results on both the mini benchmark data and a real data set.

## 2 Derivation of graph-based normalization algorithm

### 2.1 Definitions and Notations

An RNA-seq measurement on one biological sample results in a vector of abundance values of *n* genes/transcripts. A collection of measurements on *m* samples results in *m* such vectors can then be represented by an *m* × *n* matrix.

Let *A* denote the abundance matrix in which the element *a_ij_* is the true abundance of transcript *j* in sample *i*, *C* the read count matrix in which the element *c_ij_* represents the read count of transcript *j* in sample *i*. The total read counts of row *C_i_* is the sequencing depth (or library size) of sample *i*, 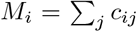. Let *A_rel_* denote the relative abundance matrix, of which the element 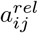 is the relative abundance of transcript *j* in sample *i*

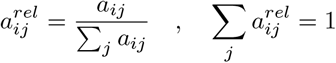

*A* is the underlying expression profile dictating the measurement in *C*. Since the exact recovery of *A* is difficult, it usually suffices to normalize *C* to a manifest abundance matrix *A*\* of which 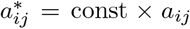, const is an unknown, yet absolute constant. Such *A*\* is considered a *desirable* normalization.

**Differential** genes/transcripts are differentially expressed due to distinct biological regulation of the conditions being studied, thus are of biological interest. The term condition may represent different states of a cell population (normal vs tumor, control vs drug-administered, etc.), or different histological origins (cerebellum vs frontal cortex, lung, liver, muscle, etc.). Although *differential* is usually encountered in the comparison of two conditions, it can be used in multi-condition assays to indicate genes/transcripts that vary with the conditions.

**Reference** genes/transcripts are expressed at equivalent levels across conditions. From the biological standpoint, reference genes/transcripts should be constitutively expressed, or at the least, are not under the biological regulations that distinguish the conditions. It should be noted that, for the purpose of normalizing read counts, reference genes/transcripts are numerically stable across conditions, and are not necessarily related to housekeeping genes/transcripts.

In this text, it is often more appropriate to use the term **feature** in place of **gene/transcript** to indicate the target entity of quantification, which can be mRNA, non-coding RNA or spike-in RNA. These features can be quantified at the transcript level or the gene level which aggregates the abundance of multiple isoforms if necessary.

### 2.2 Normalization by references

Normalization by references has been a standard practice since the early expression profiling experiments where a few transcripts are measured individually by quantitative PCR [14]. This idea has been carried over to expression profiling by high-throughput sequencing. Here we show how it works in this new setting (Theorem 1), and how the inclusion of differential features or exclusion of reference ones affect the normalization (Theorem 2,3).

In the following, 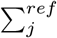 means summation over features in the reference set, and 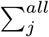 means summation over all features.

#### Theorem 1.

*Let N* = [*N*_1_, *N*_2_, …, *N_m_*]*^T^ be the reference-based normalizing vector, i.e. the scaling factor N_i_ is the sum of read counts of all the reference features in that sample*.

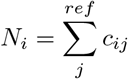

*The manifest abundance A* resulted from normalizing C against N is the desirable manifest abundance. In other words, if 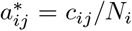 then 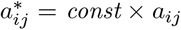, const is a quantity that does not depend on the row/column*.

**Proof.**

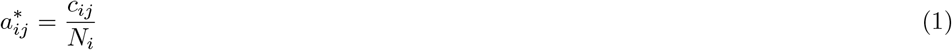

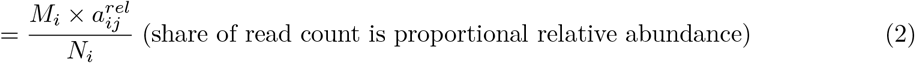

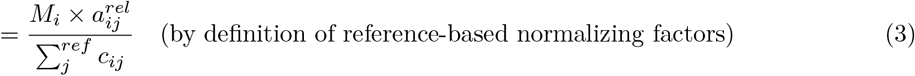

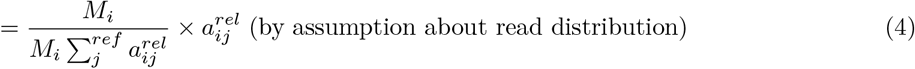

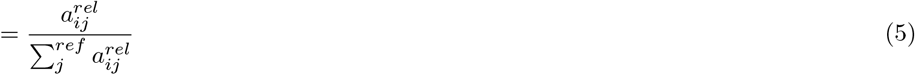

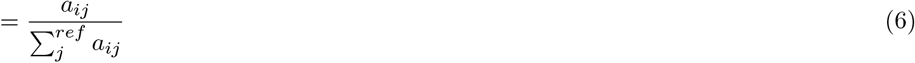

Since references are constant across samples, their sum is also constant.

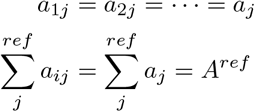

Thus, the equation (6) implies that the manifest abundance is simply a multiple of the true abundance.

#### 2.2.1 A subset of references is sufficient for normalization

Theorem 2 and 3 demonstrate the effect of mistaking differential features in the reference set, or missing some reference features during normalization.

##### Theorem 2.

*If 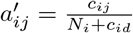 in which c_id_ is the count of a differential feature, then A′ is not a desirable normalization*.

**Proof.**

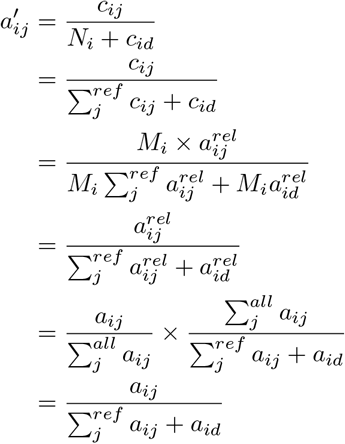

Since *d* is differential gene, the denominator is not constant, thus *A′ ≠ const × A*.

##### Theorem 3.

*Any non-empty subset of the reference set leads to a valid normalization*.

**Proof.**

Identical to that of Theorem 2.

### 2.3 Manifest correlation of transcripts in RNA-seq

In the following sections we will use *c* for read count and *T* for the true abundance, hence *c*is a function of *T*, i.e. *c = f*(*T*). The subscript *i,j* indicates different conditions, and *u, v* different features.

For simplicity, feature abundance is treated as condition-specific constant, that is, assuming biological (and technical) variance to be 0.

#### 2.3.1 *u* and *v* are both references

With this simplification, the reference features are absolute constants, i.e.

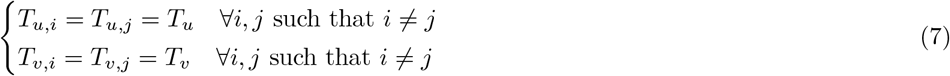

Since read counts depends on true abundance *T*, sequencing depth *M*, and transcript length *ℓ*,

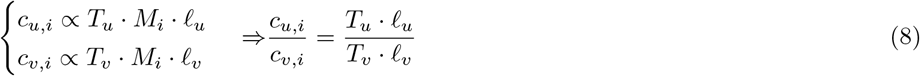

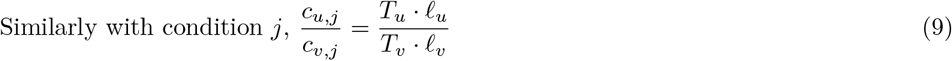

From Eq. (8) and (9), it is true that

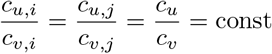

**Remark** (1). If *u* and *v* are both reference features, their read counts are linearly correlated.

#### 2.3.2 *u* is differential, *v* is reference (or vice versa)

Equivalently,

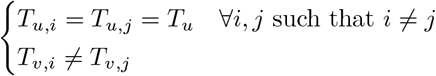

With similar operation, the observed relation between *u* and *v* in each condition are

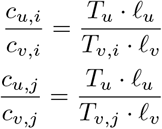

**Remark** (2). If *u* is a differential feature and *v* is a reference one (or vice versa), their read counts are not linearly correlated.

#### 2.3.3 *u* and *v* are both differential

Equivalently,

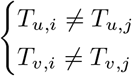

Linear correlation in this case requires that 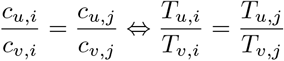

**Remark** (3). If *u* and *v* are both differential features, the two will exhibit linear correlation if and only if they vary at similar proportion (i.e. same fold change) across different conditions.

### 2.4 Graph-based normalization

It follows from the remarks in Section 2.3 that all the references in an RNA-seq data set are linearly correlated with one another. Although the derivation was based on the simplistic treatment of expression levels as condition-specific constants, the effect was in fact observed in real data (Figure 1). Using a graph to model features as vertices, and positive correlation between them as edges, it is apparent that reference features will manifest themselves as complete subgraph. However, considering Remark (3), there might exist other complete subgraphs composed of strongly co-expressed differential features. Can we distinguish the reference subgraph from the differential ones? In biological systems, such differential features must be tightly regulated and and co-expressed throughout all conditions, for example, when they are subunits of a complex which is always assembled with the same composition. Since it is less likely to encounter the coupling of very large complexes, setting a minimal size for the reference subgraph may help eliminate this mistake. For example, requiring the subgraphs to contain at least 0.1% of the features in the mouse genome is equivalent to restricting the search space to those larger than 70 genes, surpassing the 49 subunits found in the large (60S) ribosome subunit, one of the largest eukaryotic protein complexes. Among the remaining subgraphs, the most suitable ones can be selected based on the fact that all reference features are correlated, resulting in a close-to-rank-1 read count matrix. To measure this property, we used *rank-1-residuals. Rank-1-residuals* of the reference features *R*, more precisely of their read count matrix *C_R_*, is the normalized sum of the singular values except the first one.

**Figure 1.**
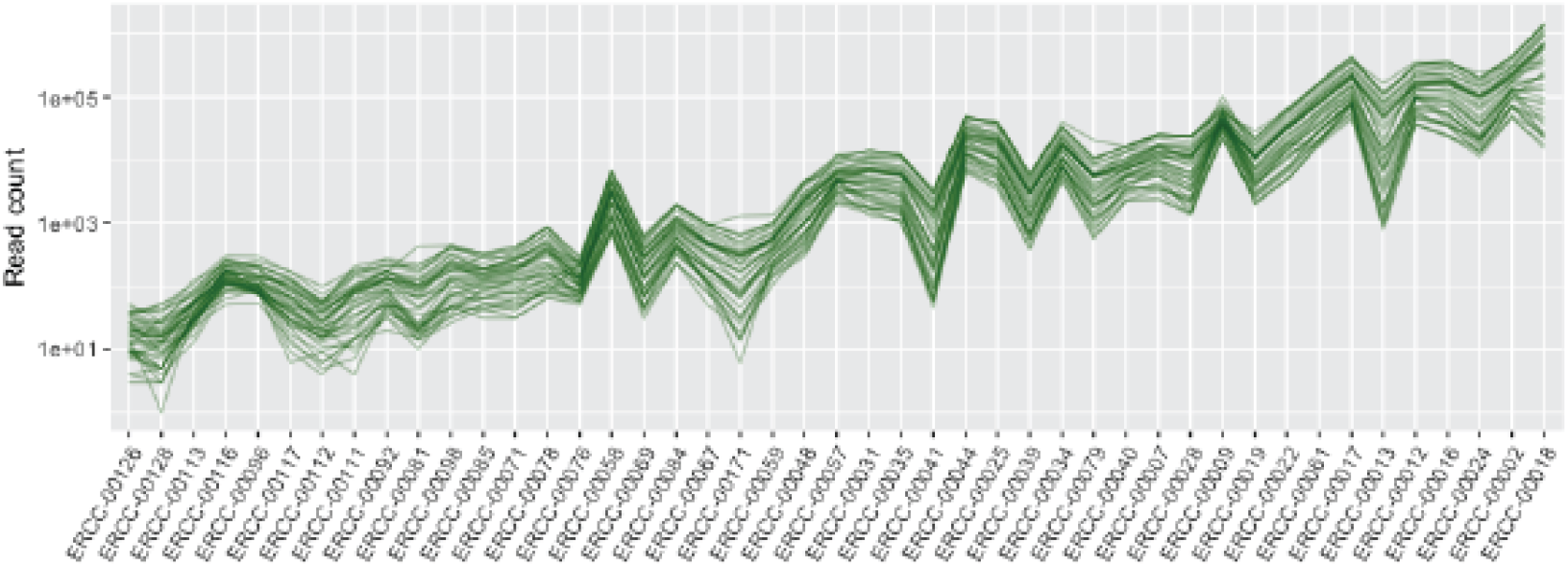
Read counts of ERCC spike RNAs in ENCODE mouse tissue samples, all of which were spiked with the same spike-in concentration. Each polyline represents a sample, spike-in RNAs are sorted by their input concentration. The parallel polylines indicate a strong correlation between these spikes.

#### Listing 1 Outline of the graph-based algorithm to identify reference features

~~~
proc identify_references (C):
    for i from 1 to n - 1
        for j from (i+1) to n
            if (cor (i,j) >= t) then *E_ij_* = 1
    G = (V,E)
    candidates = maximal_cliques (G)
    return best (candidates)
~~~

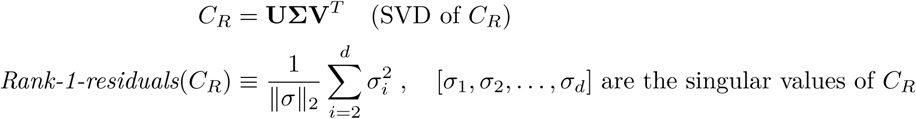

Altogether, these observations implies that by finding all `maximal_cliques()` and select the `best()` one, i.e. the lowest rank-1-residuals, one can identify the set of references (Listing 1) to be used in normalizing the read counts.

#### 2.4.1 Practical considerations

The outline in Listing 1 is in fact not efficient enough for realistic data. First, the construction of graph *G*(*V, E*) requires *O*(|*V*|^2^) both in time and space for calculating and storing the correlation matrix. A typical genome with 70000 take up 18GB, far exceeding the average computer memory of 4GB at the time of writing. The number of vertices |*V*| can be significantly reduced by retaining only features that have non-zero (or high enough to be considered reliably detected) read counts across all samples. The cumulative distribution of minimum read count across samples of the real data set revealed that 85% of the vertices can be eliminated with a non-zero filter. Second, the enumeration of all maximal cliques takes exponential time, for which the most efficient algorithm available runs in time *O*(*d*|*V*|3^*d*/3^), that is, exponential in graph degeneracy *d* which measures a graph sparsity, making it efficient only on sparse graphs where *d* is small enough [4]. To avoid this prohibitive cost, the problem is replaced by finding densely connected subgraphs which has numerous modeling approaches and corresponding solutions [7]. Since we are only concerned with one outstanding community in the graph, an accurate and complete graph partitioning may not be necessary. Furthermore, as a subset of references is sufficient for good normalization (Theorem 3), it is tolerable to miss a few members in the target community. For those reasons, any good graph partitioning method can be used in this step. We employed stochastic block model as implemented in graph-tool [11, 12].

##### Listing 2 Graph-based algorithm to identify reference features with practical considerations

~~~
proc identify_references(C):
    for i from 1 to n-1
        for j from (i+1) to n
            if (cor(i,j) >= t) then *E_ij_* = 1
    G = (V,E)
    candidates = community(G)
    remove candidates with minimum cor ≤ t
    return argmin rank-1 - residuals (b)
      *b∈candidates*
~~~

Per our visual inspection of expression data, a correlation threshold *t* ≥ 0.75 seems to indicate reasonable correlation, thus was chosen for the proof-of-concept experiments. Future studies may explore how this parameter affects the overall performance of the algorithm.

## 3 Experimental results

Data were compiled from the collection of tissue-based mouse mRNA experiments processed with the ENCODE Uniform Processing Pipeline for RNA-seq into *samples × genes* read count matrix. The full data set includes 71 samples with good enough quality (sufficient sequencing depth and read length) and 69691 genes. A subset of 41 samples that were spiked with ERCC synthetic RNAs (NIST Pool 14 concentrations) and 1837 genes (plus spike RNAs) were used to construct the benchmark data as illustrated in Figure 2B. In these data, the ERCC spike RNAs serve as reference features, while the differential features are emulated by genes known to participate in signal transduction pathways (REACTOME accession R-MMU-162582 [6]). This choice of differential genes aims to ensure that the benchmark data (1) covers a wide range of expression levels (gene products in a signaling cascade are expressed at different levels), (2) includes biologically meaningful correlation, i.e. regulated co-expression, besides artifact correlation of the reference genes and (3) mimics the variability across tissue types (signaling pathways are generally different across biological conditions). From this pool of differential features, multiple benchmark data sets are generated by sampling the combinations of signal transduction pathways (Figure 2B).

**Figure 2.**
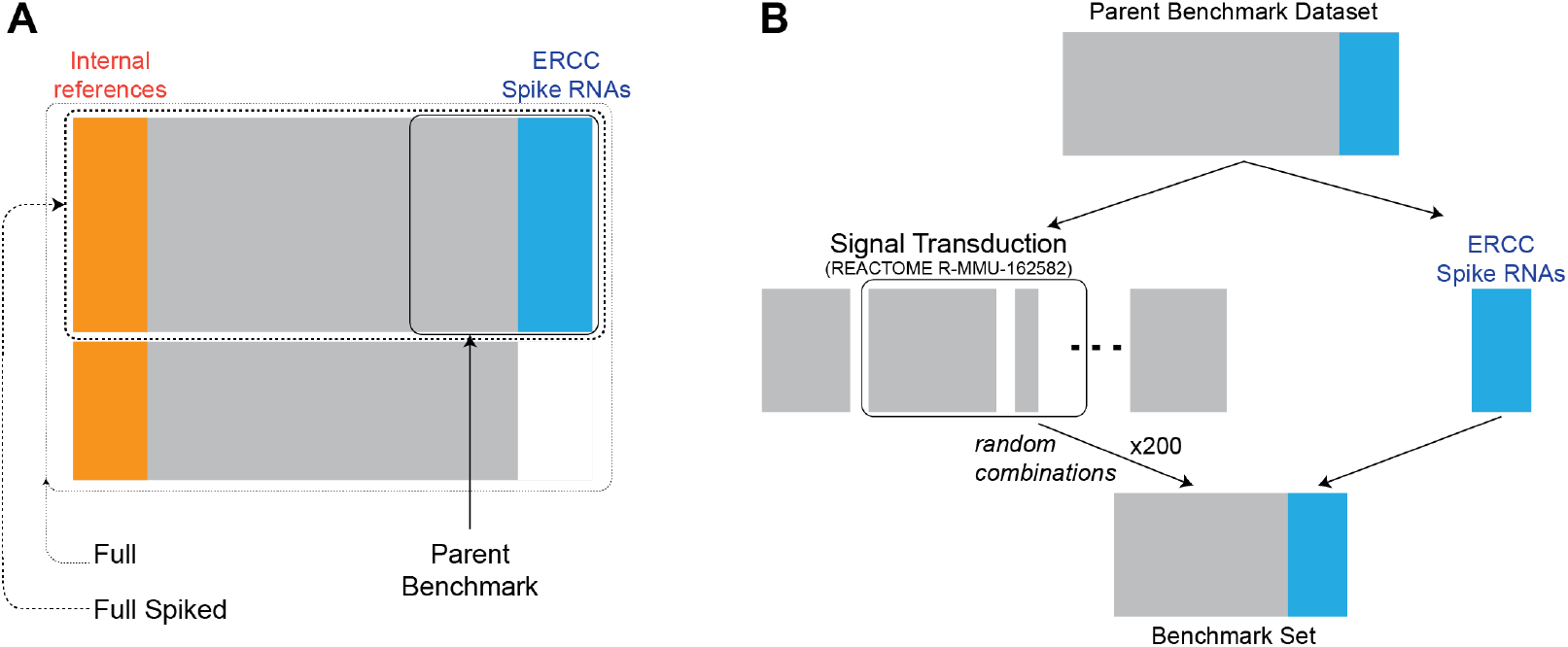
Diagram of different data sets used in this study. (**A**) The largest – *full* dataset – includes 71 samples and 69691 features. Restricting the samples to those that were spiked with ERCC spike RNAs resulted in the *full spiked* data set (41 samples × 69691 features). Restricting the genes further to those involving the signal transduction pathways resulted in the parent set that were sampled to create benchmark data. (**B**) Method of generating benchmark data sets from experimental mRNA-seq data.

### 3.1 Performance on benchmark data

It follows from the derivation that if the quantitation step has accounted for all the unwanted biases, such that read counts are distributed proportionally to relative abundance, we should observe linear correlation on the read counts. To explore the effect of correlation measures, we attempted several ways to calculate correlation, including two types of correlation measures and three different transformations on read counts (Figure 3). The good performance attained with the Pearson Correlation Coefficient on read counts (without log-transformation) implies that the quantitation tool in used has adequately accounted for those biases.

**Figure 3.**
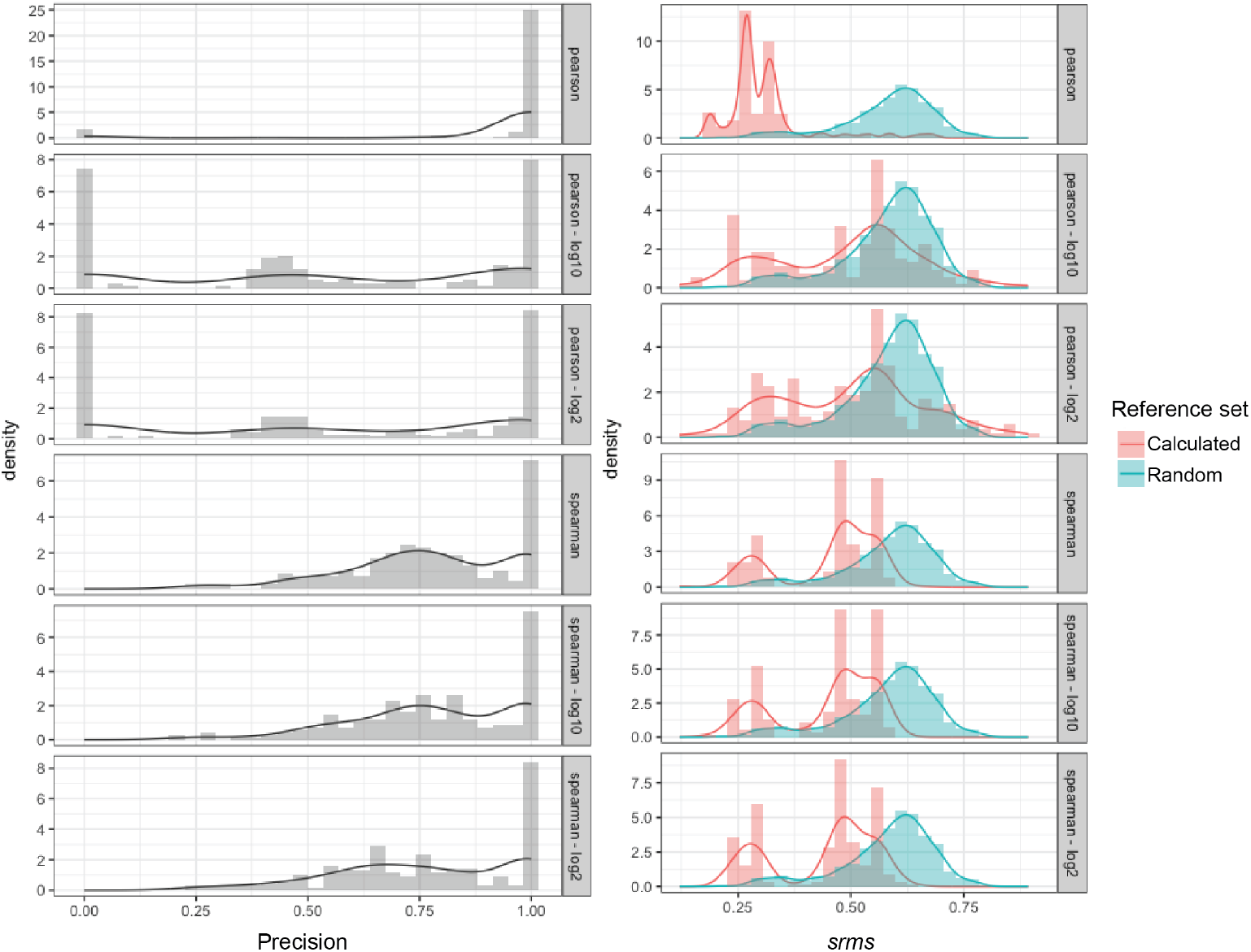
Performance of graph-based normalization on the benchmark data measured by precision in reference identification (Left) and *srms* and by deviation from the ground-truth normalization (Right). *srms* resulted from normalization against random set of references are plotted for comparison.

Performance of the normalization procedure is measured in two terms, by the *precision* in detecting the spike RNAs and by *standardized deviation (srms)* from the ground-truth normalization. Ground-truth normalization is obtained by scaling against the set of all spike RNAs. The standardized deviation of the abundance matrix *A_X_* resulted from normalizing *C* against the reference set *X* is the root-mean-squared deviation between its standardized version and that of the ground-truth. In a *standardized* abundance matrix, all features are scaled to zero mean and unit variance.

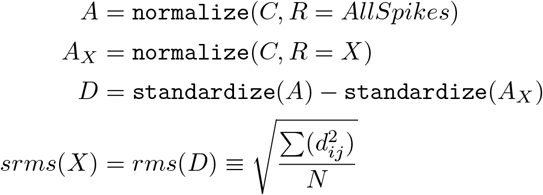

Figure 3 summarizes the performance of this algorithm on the benchmark data. The best performance was achieved with graphs built on Pearson correlation coefficients of read counts without log-transformation. In this case, one can identify the references with precision close to 1.0, and deviation from the ground-truth normalization < 0.3 (significantly smaller than that from of random reference sets) most of the time. Albeit the precision was sometimes low (close to 0.), closer examination of these cases suggested that the non-spike references identified by the algorithm are potentially valid. For example, some genes in the output reference set are in fact playing critical roles in maintaining universal living functions such as *Ccnt2* (cyclin T2), *Polr2d* (RNA polymerase II polypeptide D), *Polr2l* (RNA polymerase II peptide L), *H3f3b* (H3 histone family 3B), *Cenpc1* (centromere protein). It may be necessary to have a more carefully curated list of genes emulating differential features in the benchmark data set for more accurate evaluation.

In comparison with a state-of-the-art method, TMM [13], *srms* deviation of graph-based normalization (gbnorm) is always lower, and the distributions of *srms* deviation showed a distinctively better performance on the benchmark data.

### 3.2 Performance on real data

The best setting of the above procedure, i.e. graph built with PCC on read counts without any transformation, was applied on the full data set of 71 samples × 69691 genes, resulting in 23 references. If these genes are references is the full set, they are also references in any subset of samples and can be used to normalize these subsets. To evaluate the quality of this method on the real data, we used the full spiked set which has both internal and external references (Figure 2A). The abundance matrix obtained by normalizing against by external (spike) references serves as the ground truth normalization. The internal reference set determined as above were used to scale the full spike count matrix, resulting in the graph-based normalization. TMM was also used to normalize the same set. As shown in the table below, graph-based normalization resulted in a better (smaller) *srms* compared to TMM. Although the graph-based method took much longer, approximately 43 minutes, normalization is generally a one-time operation, rendering this running time reasonably accessible. Since this processing step critically affects all downstream analyses and biological interpretations, a better normalization should always be preferred if it is accessible.

**Figure 4.**
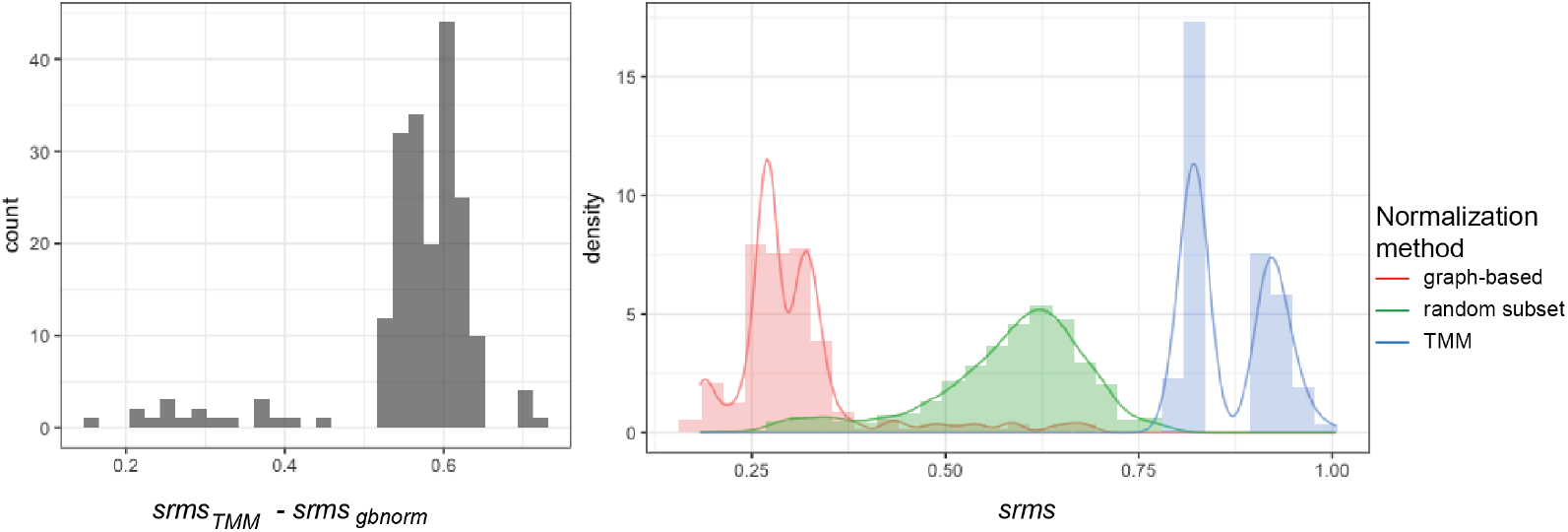
Comparison of graph-based normalization and TMM, a state-of-the-art method, on benchmark data. *Left* – Discrepancies between TMM-normalized and graph-based normalized expression levels, measured in *srms* against the ground-truth normalization. *Right* – Distribution of *srms* deviation from the ground-truth normalization.

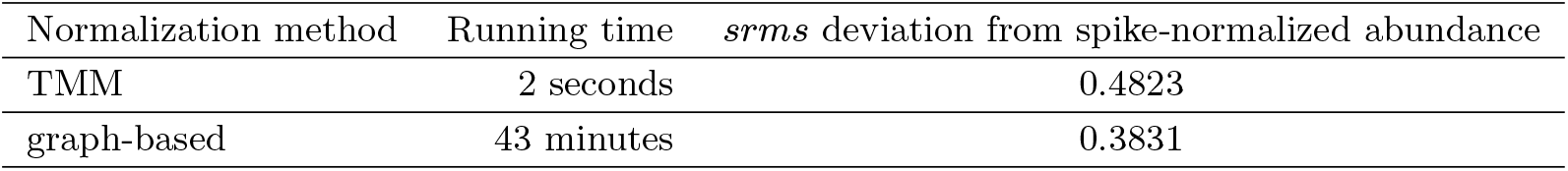

| Normalization method | Running time | $srms$ deviation from spike-normalized abundance |
| --- | --- | --- |
| TMM | 2 seconds | 0.4823 |
| graph-based | 43 minutes | 0.3831 |

## 4 Conclusions and future directions

We proposed a new method to normalize RNA-seq data. Unlike the existing methods, this algorithm helps identify a set of internal references based on their correlation in read counts, thus eliminating the need for prior knowledge about stably expressed genes, assumptions about experimental conditions, and external spike-ins. It worths noticing that this method requires a large number of samples and a quality feature count method for correlation measure to be reliable. That said, these requirements are generally indispensable for any RNA-seq processing workflow. The algorithm involves several parameters, including the choice of correlation measure, the correlation threshold to form an edge, the ranking measure of candidate reference set, the community detection algorithm to be used and its corresponding parameters. Future work may explore how each parameter affect the performance of reference identification as well as normalization. Beyond these parameters, it is also important to understand how the method perform in various practical settings, specifically gene-level *vs* transcript-level read counts, the degree of heterogeneity among conditions, the proportion of differential features. Another important task is to compare the performance of this new method against the others. Since evaluation depends on the simulation data, care should be taken to generate benchmark data sets that are as realistic as possible.

## Footnotes

1 *Normalization by library size* is not considered between-sample normalization.

## Acknowledgements

We want to thank Dr Jay Gertz for insightful discussions, Dr Jeff Phillips for helping to improve the manuscript, and Jie Shi Chua for proofreading.

## References

1 Chien-Ming Chen, Yu-Lun Lu, Chi-Pong Sio, Guan-Chung Wu, Wen-Shyong Tzou, and Tun-Wen Pai.Gene Ontology based housekeeping gene selection for RNA-seq normalization. Methods, 67(3):354–363, June 2014. doi:10.1016/j.ymeth.2014.01.019.

2 Kaifu Chen, Zheng Hu, Zheng Xia, Dongyu Zhao, Wei Li, and Jessica K. Tyler. The Overlooked Fact: Fundamental Need for Spike-In Control for Virtually All Genome-Wide Analyses. Molecular and Cellular Biology, 36(5):662–667, January 2016. doi:10.1128/ MCB.00970-14.

3 Marie-Agnès Dillies, Andrea Rau, Julie Aubert, Christelle Hennequet-Antier, Marine Jeanmougin, Nicolas Servant, Céline Keime, Guillemette Marot, David Castel, Jordi Estelle, Gregory Guernec, Bernd Jagla, Luc Jouneau, Denis Laloë, Caroline Le Gall, Brigitte Schaёffer, Stéphane Le Crom, Mickaël Guedj, and Florence Jaffrézic. A comprehensive evaluation of normalization methods for Illumina high-throughput RNA sequencing data analysis. Briefings in Bioinformatics, 14(6):671–683, January 2013. doi:10.1093/bib/bbs046.

4 David Eppstein,Maarten Löffler, and Darren Strash.Listing All Maximal Cliques in Sparse Graphs in Near-optimal Time. *arXiv:1006.5440 [cs]*, June 2010. arXiv:1006.5440.

5 Ciaran Evans, Johanna Hardin, and Daniel M. Stoebel. Selecting between-sample RNA-Seq normalization methods from the perspective of their assumptions. Briefings in Bioinformatics, February 2017. doi:10.1093/bib/bbx008.

6 Antonio Fabregat, Steven Jupe, Lisa Matthews, Konstantinos Sidiropoulos, Marc Gillespie, Phani Garapati, Robin Haw, Bijay Jassal, Florian Korninger, Bruce May, Marija Milacic, Corina Duenas Roca, Karen Rothfels, Cristoffer Sevilla, Veronica Shamovsky, Solomon Shorser, Thawfeek Varusai, Guilherme Viteri, Joel Weiser, Guanming Wu, Lincoln Stein, Henning Hermjakob, and Peter D’Eustachio. The Reactome Pathway Knowledgebase. Nucleic Acids Research, 46(D1):D649–D655, January 2018. doi:10.1093/nar/gkx1132.

7 Santo Fortunato and Darko Hric. Community detection in networks: A user guide. Physics Reports, 659:1–44, 2016.

8 J. Huggett, K. Dheda, S. Bustin, and A. Zumla. Real-time RT-PCR normalisation; strategies and considerations. Genes and Immunity, 6(4):279–284, June 2005. doi: 10.1038/sj.gene.6364190.

9 Lichun Jiang, Felix Schlesinger, Carrie A. Davis, Yu Zhang, Renhua Li, Marc Salit, Thomas R. Gingeras, and Brian Oliver. Synthetic spike-in standards for RNA-seq experiments. Genome Research, 21(9):1543–1551, January 2011. doi:10.1101/gr.121095.111.

10 Yanzhu Lin, Kseniya Golovnina, Zhen-Xia Chen, Hang Noh Lee, Yazmin L. Serrano Negron, Hina Sultana, Brian Oliver, and Susan T. Harbison. Comparison of normalization and differential expression analyses using RNA-Seq data from 726 individual Drosophila melanogaster. BMC Genomics, 17, January 2016. doi:10.1186/s12864-015-2353-z.

11 Tiago P. Peixoto. The graph-tool python library, May 2017 doi:10.6084/m9.figshare.1164194.v14.

12 Tiago P. Peixoto. Nonparametric Bayesian inference of the microcanonical stochastic block model. Physical Review E, 95(1):012317, January 2017. doi:10.1103/PhysRevE. 95.012317.

13 Mark D. Robinson and Alicia Oshlack. A fffor differential expression analysis of RNA-seq data. Genome Biology, 11:R25, 2010. doi:10.1186/ gb-2010-11-3-r25.

14 Jo Vandesompele,Katleen De Preter,Filip Pattyn, Bruce Poppe, Nadine Van Roy, Anne De Paepe, and Frank Speleman. Accurate normalization of real-time quantitative RTPCR data by geometric averaging of multiple internal control genes.Genome biology, 3(7):research0034–1,2002.

15 Zhong Wang, Mark Gerstein, and Michael Snyder. RNA-Seq: A revolutionary tool for transcriptomics. Nature Reviews Genetics, 10(1):57–63, January 2009. doi:10.1038/nrg2484.

16 Bin Zhuo, Sarah Emerson, Jeff H. Chang, and Yanming Di. Identifying stably expressed genes from multiple RNA-Seq data sets. PeerJ, 4, December 2016. doi:10.7717/peerj.2791.

